# Adaptive management plans rooted in quantitative ecological predictions of ecosystem processes: putting monitoring data to practical use

**DOI:** 10.1101/2020.10.11.334789

**Authors:** Christian Damgaard

## Abstract

It is demonstrated how a hierarchical structural equation model that is fitted to temporal ecological monitoring data from a number of sites may be used to generate local ecological predictions and how these local ecological predictions may form the basis of adaptive management plans. Local ecological predictions will be made for the cover of cross-leaved heath on Danish wet heathlands, which is one of the indicators that determine the conservation status of wet heathlands under different management scenarios. Based on a realistic example, the model predictions concludes that grazing by domestic herbivores on wet heathlands with a relatively low cover cross-leaved heath cannot be recommended as the only management practice. Generally, it is recommended to use ecological monitoring data to generate quantitative and credible local adaptive management plans where the uncertainty is taken into account.

## Introduction

Anthropogenic environmental changes, e.g. land-use and climate changes, put pressures on biodiversity in natural and semi-natural ecosystems (Dawson et al. 2011; IPBES 2019; Jung et al. 2019; Timmermann et al. 2015). If ecosystems are at least partly controllable, i.e. management actions may have positive effects on biodiversity, then a possible societal response to environmental pressures on biodiversity is to manage ecosystems in order to achieve specified conservation goals. However, due to complicated ecosystem dynamics with considerable time-lags, the effect of management actions on biodiversity are often uncertain and it has been recommended to implement adaptive management plans, where our uncertainties on the ecological processes are taken into account (Abrahms et al. 2017; Williams et al. 2009).

Adaptive management is a structured decision making process especially targeted the management of natural and semi-natural ecosystems when a number of requirements are fulfilled (Table 1). The concept of adaptive management is defined by Williams et al. (2009): “… as a decision process that promotes flexible decision making that can be adjusted in the face of uncertainties as outcomes from management actions and other events become better understood. Careful monitoring of these outcomes both advances scientific understanding and helps adjust policies or operations as part of an iterative learning process. … .”

**Table 1.**
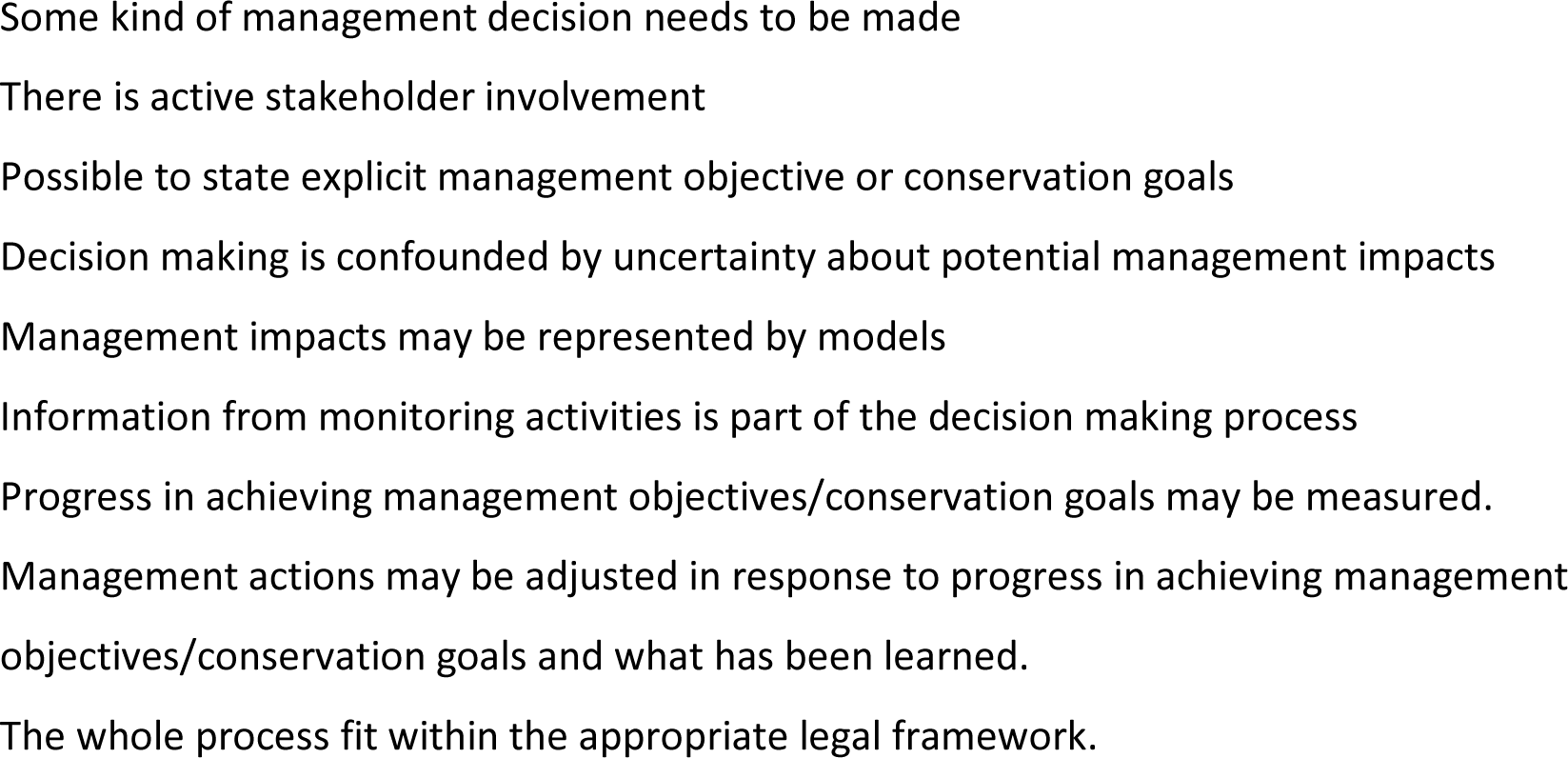
Requirements that should be fulfilled for adaptive management to be an appropriate method of decision making (adapted after Williams et al. 2009).

The development of predictive ecological models that incorporates the different sources of uncertainty is a key feature of the adaptive management process. Although such modelling efforts in specific ecosystems at first sight may seem premature due to lack of data or system knowledge, it will usually be valuable to incorporate the existing system knowledge in some modelling framework. As stated by Williams et al. (2009): “A common complaint used to justify not undertaking an adaptive program is that the data are sparse and there is too much uncertainty to build models. But this is precisely where adaptive management is most valuable—in expressing and reducing uncertainty. The alternative to building models of system dynamics is to allow the assumptions of decision makers and stakeholders— essentially, the models that exist in the minds of a few individuals—to remain unexpressed and untested.”

Currently, most ecological predictions are constructed by modelling observed ecological patterns, i.e. by modelling spatial variation among environmental and ecological variables without considering the underlying ecological processes. In such spatial regression models, it is implicitly assumed that the observed spatial patterns are due to different realizations of the equilibrium properties of the studied ecological process and that the temporal dynamics at the site level may be ignored. This space-for-time approach is often problematic (Damgaard 2019a), and here, the ecological predictions are based on estimated temporal dynamics, which are expected to encapsulate the underlying ecological processes. This approach is possible because we have access to comprehensive environmental and ecological monitoring data in both space and time (Nielsen et al. 2012), where it is possible to statistically separate the observed temporal effects from the spatial variation. Furthermore, by using a Bayesian hierarchical framework it is possible to separate different sources of measurement- and sampling errors from the process error and thereby get a more precise estimate of the process uncertainty, which is needed for generating ecological predictions and adaptive management plans (Damgaard 2019b).

In this study it is demonstrated how a hierarchical structural equation model that is fitted to temporal ecological monitoring data from a number of sites may be used to generate local ecological predictions and how these local ecological predictions may form the basis of adaptive management plans. The approach is generic, but will be exemplified using wet heathlands, which are relatively species-poor, semi-natural habitats at sandy nutrient-poor soils, where the water table is naturally high and the vegetation is mainly comprised of dwarf shrubs and graminoids (Hampton 2008). The dwarf shrub cross-leaved heath (*Erica tetralix*), which is a characteristic plant species for the habitat, has experienced a general decrease in cover over the past 15 years (Damgaard 2012; Damgaard et al. 2017). Furthermore, the cover of cross-leaved heath is one of the indicators that determine the conservation status of wet heathlands (high conservation status: > 0.3, low conservation status: < 0.1) in the Danish multi-criteria assessment system (Damgaard et al. 2019; Nygaard et al. 2014), which is used for reporting the habitat conservation status under the EU habitat directive (EU 1992).

## Materials and methods

### Ecological and environmental data

Cover data of all higher plant species from 441 plots with wet heathland vegetation at 39 Danish sites were sampled irregularly in the time-period from 2007 to 2014 using the pin-point method, where the square frame (50 cm X 50 cm) consisted of 16 grid points that were equally spaced by 10 cm (Nielsen et al. 2012). The plots were resampled with GPS-certainty (< 10 meters), and including resampling there was cover data from 1322 plots. For model simplification purposes, the plant species were classified into the three key dominating species of wet heathlands (*Erica tetralix, Calluna vulgaris*, and *Molinia caerulea*) and an aggregated class of all other higher plant species (Damgaard 2019b).

The environmental variables that were assumed to be most important for plant community dynamics in wet heathlands *a pri*ori were selected under the constraint that the data were available at a relevant resolution (Table 2). Note that it was not possible to obtain data on the distance to the water table and its seasonal fluctuations, which may be an important factor in regulating the vegetation at wet heathlands.

**Table 2.**
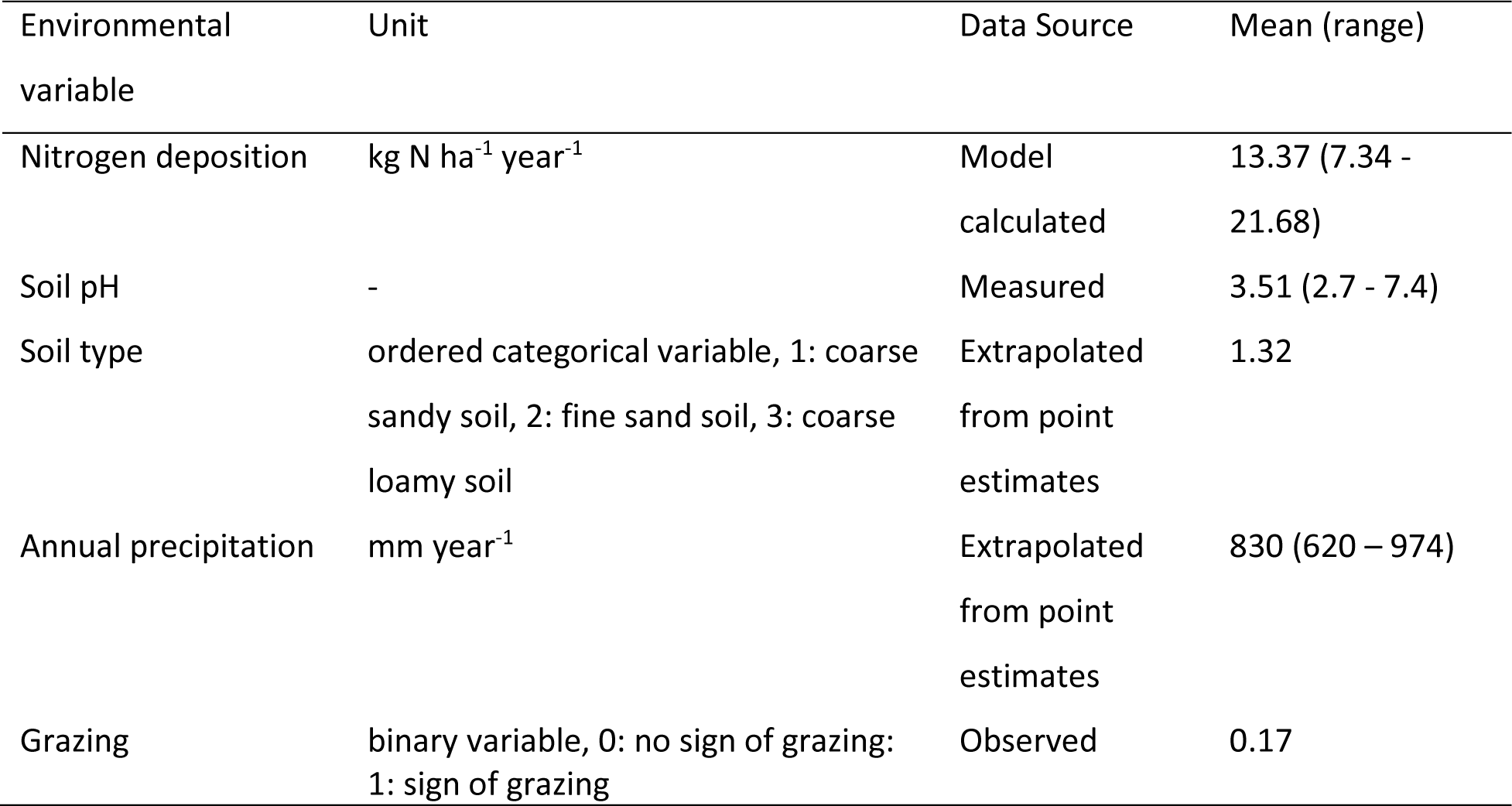
Selected environmental variables, units, data sources and their mean values and ranges in Danish wet heathlands.

### Hierarchical structural equation model of wet heathlands

The used hierarchical structural equation model (SEM) and the results when the model is applied to the Danish wet heathland monitoring data have been explained in detail in a previous study (Damgaard 2019b) and will only briefly be summarized here. The selected environmental variables were modelled in a SEM based on the current knowledge on the causal effect relationships among variables and their effect on the vegetation in wet heathlands (Damgaard et al. 2017; Damgaard et al. 2014; Damgaard et al. 2013; Lykke et al. 2015; Strandberg et al. 2012). Additionally, geographic regional latent effects were modelled after the Danish wet heathland sites were divided into seven geographic regions. The SEM was fitted within a Bayesian hierarchical framework with structural equations and measurement equations in an acyclic directed graph (Fig. 1), where spatial and temporal processes are separated and each arrow is modelled by a conditional likelihood function (Damgaard 2019b).

**Fig. 1.**
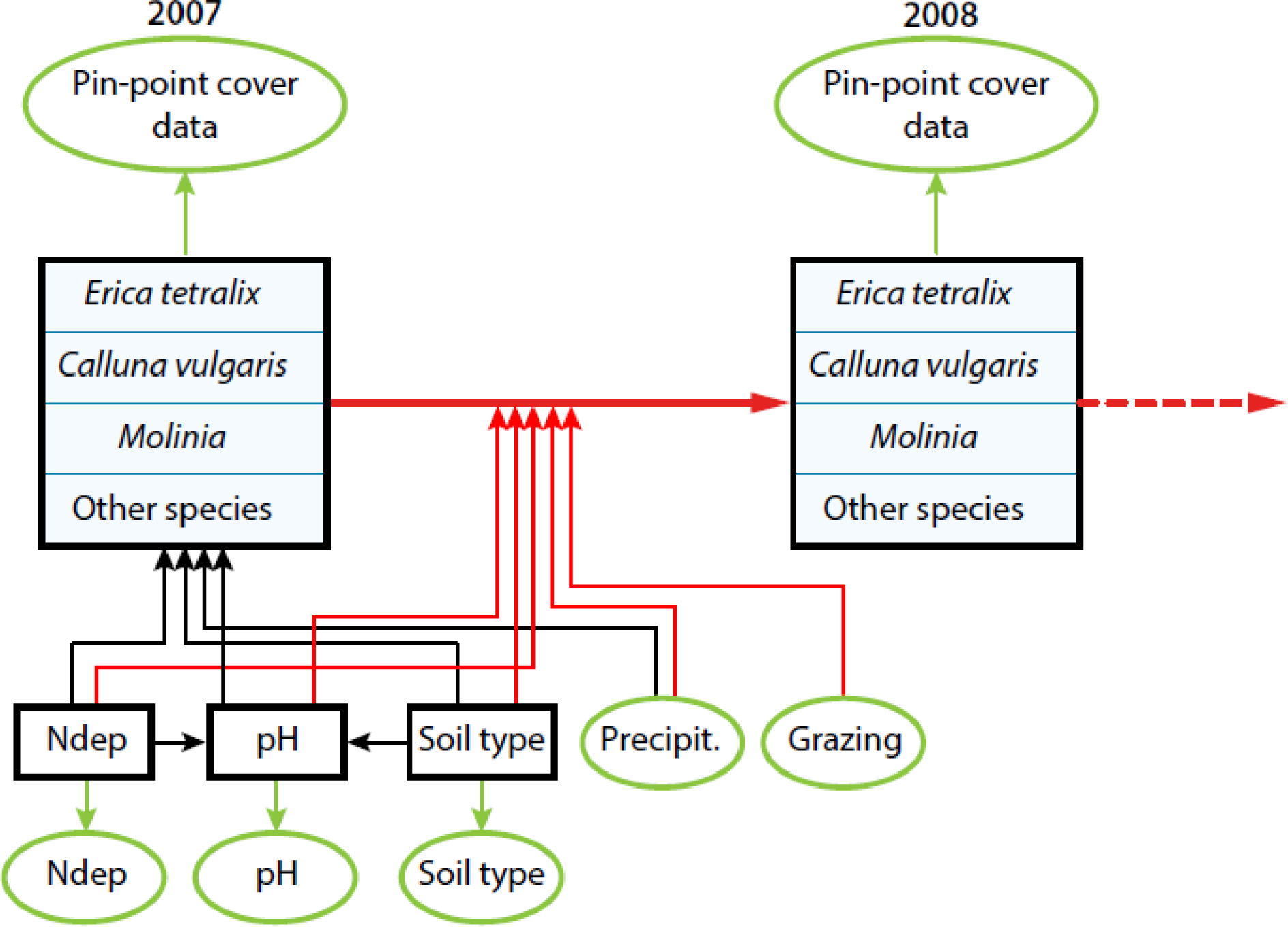
Outline of the structural equation model. The spatial variation in vegetation cover in 2007 is modelled by nitrogen deposition (Ndep), soil pH (pH), soil type and precipitation (Precipit.). The yearly change in vegetation cover from 2007 to 2014 (only a single yearly change is shown in the figure) is modelled by all the former variables as well as grazing. The black boxes are latent variables and the green ovals are data. The black arrows denote spatial processes, and the red arrows denote temporal processes (Damgaard 2019b).

The resulting joint posterior probability distribution of model parameters allowed us to test various general ecological hypotheses on how wet heathland vegetation is regulated as well as quantifying the effects of the selected environmental variables on the observed changes in the vegetation. Overall, the temporal model explained the observed changes in plant cover data rather well, and better than the spatial model explained the spatial variation in plant cover data (Damgaard 2019b). The fitted model showed that the cover of the two dwarf shrubs, bell heather and heather (*Calluna vulgaris*), increased with relatively high nitrogen deposition, high pH, sandy soil, low rainfall and absence of grazing. Nitrogen deposition had a non-significant effect on soil pH, but sandy soils had a lower pH. This means that there was both a direct effect of a relatively sandy soil on the vegetation and an indirect effect mediated via a lower pH (Damgaard 2019b).

The fitted model is a statistical model that summarizes trends and variation in the collected monitoring data and the observed correlations are not necessarily supported by current ecological working hypotheses. For example, it was surprising that nitrogen deposition on average had a positive effect on the coverage of dwarf shrubs, and this may be explained by possible covariation with other unmeasured important factors (Damgaard 2019b).

### Local ecological predictions and adaptive management plans

Using the monitoring data and the causal relationships hypothesized in the SEM, it is now possible to generate ecological predictions at a local site. The globally fitted information on the causal relationships at all the monitored sites is contained in the joint posterior probability distribution of the model parameters. Therefore, the probability distribution of the modelled ecological variable at the local site at some time in the future may be calculated from the joint posterior probability distribution by inserting local values of the ecological and environmental variables into the temporal SEM model.

Such local ecological predictions of relevant ecological variables may be used to generate adaptive management plans. For example, suppose that the conservation goal in a nature conservation project of a wet heathland site is to increase the cover of bell heather from 5% to 10% over a five-year period. Then it will be relevant to calculate the probability distribution of observing such an increase in the cover of bell heather under different management scenarios. The calculated probability distributions of the expected change in the cover of bell heather will i) help the local manager to select among different management strategies, such that the management at the site is in agreement with the conservation goals, and ii) help to set threshold levels in the adaptive management plan where the local management strategy should be reconsidered. For example, if the cover of bell heather drops significantly below the initial 5% and model predictions have shown such a change in cover to be unlikely, then the dynamics at the local site are not in agreement with the majority of the monitored wet heathland sites and the manager must consider alternative management strategies.

In order to insert local values of the ecological and environmental variables into the temporal SEM model some of these variables has to be measured at the local site. If the local ecological and environmental measurements are based on a sample of ca. 40 plots, this should be sufficient to obtain the required precision for making credible ecological predictions. Compared to the cost of the different management practices, it is relatively inexpensive to collect the necessary environmental and ecological data, and, in addition, the collected data can be used to document the effect of the management. In the following ecological predictions, it is assumed that the values of the environmental variables are known with high precision. However, any measurement- and sampling errors of the ecological and environmental variables may be incorporated into the ecological predictions using standard methods (Carroll et al. 2006).

All calculations are made using *Mathematica* (Wolfram 2020), and the relevant *Mathematica* notebook may be obtained upon request.

## Results

The probability distribution of the predicted cover of bell heather after five years at a hypothetical wet heathland site, where we want to make ecological predictions, was calculated at different scenarios, assuming that the initial cover of bell heather was 5% (Fig. 2). Here four scenarios are investigated: with and without grazing by domestic herbivores at two different soil pH. The values of the other ecological and selected environmental variables were set within a realistic range for Danish conditions, i.e. nitrogen deposition, soil and precipitation were assumed to be 15 kg N ha^−1^ year^−1^, coarse-sand soil, 700 mm year^−1^, respectively. The initial cover of *C. vulgaris, M. caerulea* and the aggregated class of all other higher plant species was set to 30%, 30%, 35%, respectively. The geographic region of the site was not specified, i.e. there were no geographic regional latent effects.

**Fig. 2.**
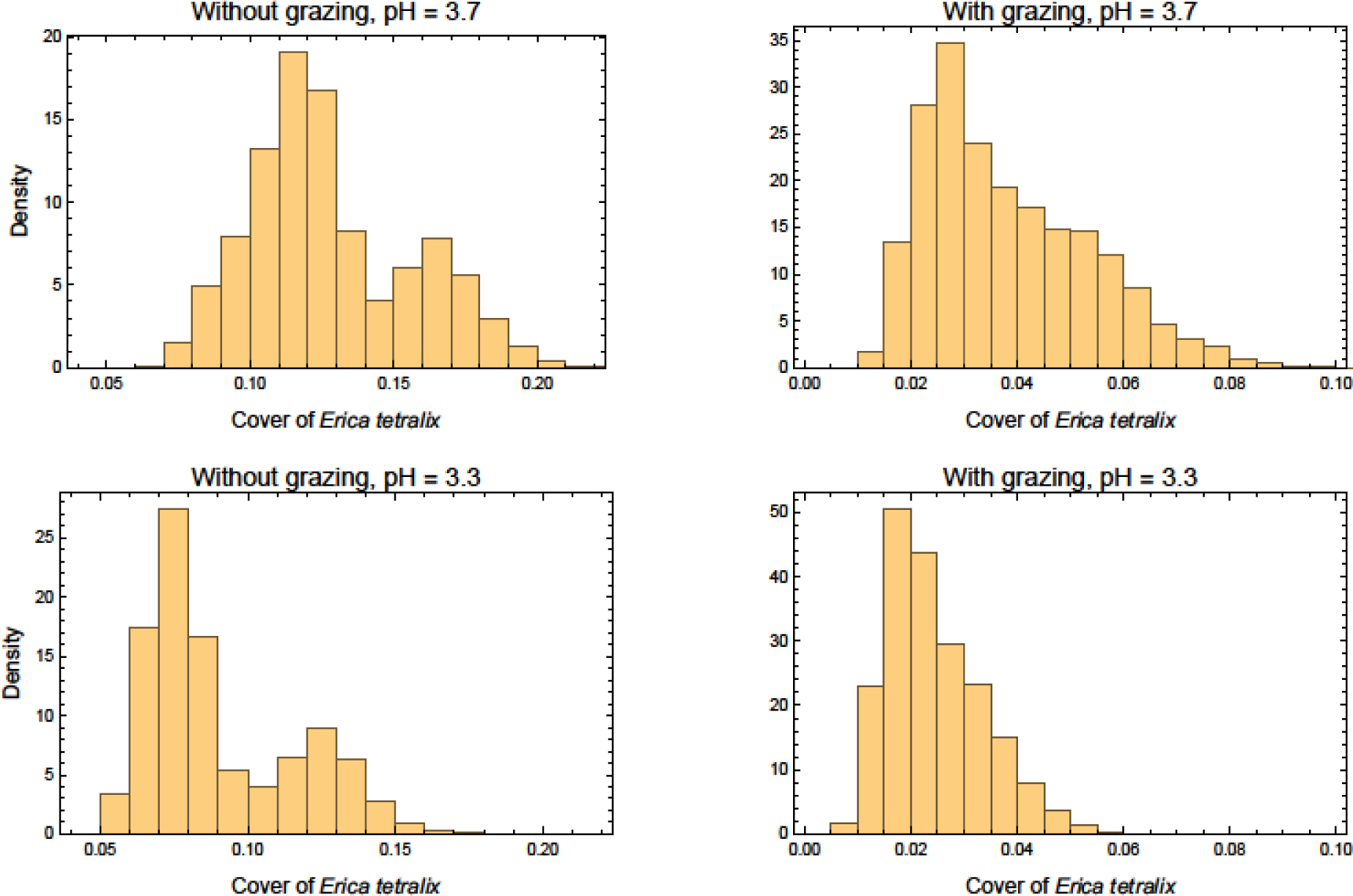
Probability densities of the predicted cover of bell heather (*Erica tetralix*) after five years with an initial cover of 5%. Four scenarios were examined: with and without grazing at two levels of soil pH. The local nitrogen deposition, soil and precipitation are assumed to be 15 kg N ha^−1^ year^−1^, coarse-sand soil, 700 mm year^−1^, respectively.

If the conservation goal at a wet heathland site was set to 10% cover of bell heather after five years, then the probability of achieving this goal depends on both soil pH at the site and whether grazing is part of the management practice. If the soil pH at the site is relatively high, the probability of reaching the conservation goal increases, whereas grazing by domestic herbivores is predicted to have an important negative effect on the probability of achieving the conservation goal (Fig. 2). Unfortunately, the grazing data only consists of a binary recording of whether there were signs of grazing by domestic herbivores, and we do not have data on which domestic herbivores were used to graze the site, nor do we know their stocking density or at which time period the sites were grazed. Therefore, the estimated effect of grazing on the vegetation at wet heathland sites is only credible when grazing by domestic herbivores is performed in a way that is typical for the sites where the monitoring data have been collected. In Denmark, grazing by domestic herbivores at wet heathland sites mainly occurs in the plant growth period from Maj to October. The most commonly used species of domestic herbivore are sheep, although cows or ponies also may be used. The recommended stocking density of large herbivores for Danish wet heathlands is less than 0.5 herbivore equivalents per hectare, where an adult sheep counts as 0.15 of a large herbivore (Miljøstyrelsen). Consequently, the model predicted effect of grazing is expected to be credible for summer grazing by sheep at the recommended stocking rate, whereas e.g. the predicted effect of whole-year grazing by ponies is less credible.

In this demonstration of the method, only soil pH and grazing varied, but if any of the other environmental variables at the hypothetical local site had varied, this would also affect the probability of achieving the conservation goal (results not shown).

Since the probability of reaching the conservation goal is calculated as probability densities, the uncertainty of reaching the goal is automatically calculated and shown in a transparent way. The source of this uncertainty is mainly variation in the temporal processes among the monitored 39 wet heathland sites, and it is expected that this number of sites is sufficiently high to contain most of the variation in the studied temporal processes at Danish wet heathland sites.

The actual adaptive management plan will depend on local conditions, stakeholders and previous experience of different management actions on wet heathlands (Williams et al. 2009). However, according to the fitted temporal SEM model it is highly unlikely that the conservation goal of 10% cover of bell heather would be reached if the local site is grazed by domestic herbivores, so it would not be realistic or credible to suggest grazing by domestic herbivores as the only management practice in the adaptive management plan. Furthermore, if the cover of bell heather in the absence of grazing decreases to a threshold level specified in the adaptive management plan, e.g. 3%, then it may be concluded that the local dynamics are atypical, and the adaptive management plan must contain alternative management actions for this case, e.g. cutting and burning (Hampton 2008). In all cases, it will be important to document the effect of the management for future learning and more specific ecological modelling of the effect of different management practices on wet heathlands.

## Discussion

Anthropogenic environmental changes are at the basis of the current worldwide biodiversity crisis (IPBES 2019), and one possible societal response to the observed decline in biodiversity is to actively manage threatened natural and semi-natural habitats. More specifically, EU member states are legally obligated to ensure that the area of each habitat type with favorable conservation status does not decrease (EU 1992). In order to prioritize and maximize the effect of management efforts of habitats, we need to be able to generate adaptive management plans that are rooted in quantitative local ecological predictions where the uncertainties are taken into account (Abrahms et al. 2017; Williams et al. 2009). The calculated ecological predictions are important in the applied management work of specifying realistic concrete conservation goals and formulating credible adaptive management plans. Such adaptive management plans, which may be assessed and evaluated by the competent authorities, may also lead to higher societal investments in biodiversity management, since the possible benefits and risks of the investment have been quantified.

In this demonstration study, I have focused on the effect of grazing on wet heathlands that differ in soil pH. As demonstrated here and in Damgaard (2019b), grazing by large domestic herbivores is expected to reduce the cover of cross-leaved heath, which is a habitat-characteristic plant species and is one of the ecological indicators that determine the conservation status of wet heathlands in the Danish multi-criteria assessment system (Damgaard et al. 2019; Nygaard et al. 2014). Consequently, based on these results, grazing by domestic herbivores on wet heathlands with a relatively low cover cross-leaved heath cannot be recommended as the only management practice.

The presented approach is generic, and different types of ecological monitoring data may be used to generate local ecological predictions and adaptive management plans. More specifically, the presented SEM is currently being fitted to monitoring data of other light open habitat types, e.g. acid grasslands and dry heathlands.

Since the uncertainties of the ecological predictions are quantified by probability distributions, the ecological predictions will also suggest levels of the ecological variables that may act as early warning in the adaptive management plans for changing the management at the site. For example, if the cover of bell heather in an ungrazed wet heathland site decreases from an initial of 5%, then the development at the site may be reasoned to be atypical and alternative measures need to be considered.

For some management practices, there may be important time lags between the onset of the management practice and when the effect of the management may be observed in the ecosystem. Such time lags are usually not contained in the ecological predictions and must be considered and incorporated into the adaptive management plans.

Some environmental data that are important for the regulation of the studied ecosystem may not be available, e.g. distance to water table data or better grazing data. Such unmeasured or poorly measured variables will surely increase the uncertainties of the ecological predictions compared to a situation where the data are available and the variable included in the SEM model. However, if the unmeasured variables at the local site are within the domain of the unmeasured variables of the sites that are monitored, i.e. the local site is not atypical, then the local ecological predictions most likely will be unbiased. We may criticize statistical models for lack of precision and not including all relevant information. However, we will not get sufficient data to make absolute precise local ecological predictions in our lifetime and, perhaps, never. If we cannot obtain absolute precision, then the next best option is to quantify our uncertainty.

In Denmark, the ecological monitoring is collected as part of the national responsibility outlined in the EU habitat directive (EU 1992) and is primarily used to report on the conservation status of the different terrestrial habitats. Why not use the often-excellent ecological monitoring data, which are already being collected in many countries, and put the global learning available in ecosystem modelling of the monitored sites to practical use by generating quantitative and credible local adaptive management plans? Next, is the current generation of applied ecologist and local managers of natural habitats ready to embrace the quantitative methods of ecological predictions?

